# Restoring declining species through translocations: A test case using flightless grasshoppers in an urban setting

**DOI:** 10.1101/2023.05.29.542782

**Authors:** Hiromi Yagui, Michael R. Kearney, Ary A. Hoffmann

## Abstract

1. Population translocations are used increasingly as a conservation strategy for vertebrates. However, relatively few attempts have been made to translocate invertebrates despite their potential benefits for food webs, and despite the practicality of undertaking such translocations in small areas including urban environments where space is limited.
2. We conducted 36 translocations of 1851 individuals of the generalist flightless grasshopper *Vandiemenella viatica* across urban Melbourne, where 93% of its original habitat has been lost.
3. We aimed to understand characteristics essential for grasshopper persistence and to investigate detection, occupancy, dispersal, and habitat suitability throughout its active period to improve insect translocation success in urban settings using revegetated sites and small remnant habitats. We also measured movement and detection probability after one week in short-term trials.
4. The one-week trial indicated that grasshopper sex and colour morph did not influence the probability of detection, and there was no evidence of directional movement by females and males.
5. One year after translocation, *V. viatica* were found in 28 out of 36 translocation sites. These surveys showed that detection probability changed across survey seasons and was influenced by soil temperature. Also, soil temperature positively affected detection in the winter surveys. Occupancy probability was influenced by graminoid cover, plant species richness and weed cover. We found no evidence of directional movement by females and males in the F1 generation. Abundance and presence/absence data were best explained by graminoid cover and plant species richness.
6. Our findings suggest that wingless grasshopper translocations are feasible in small urban patches of suitable habitat, helping to restore invertebrate biodiversity and ecological services.

## 1. Introduction

Conservation translocations are human-mediated movements of organisms from one place to another to improve a species’ conservation status and/or recuperate natural ecological processes and functions (IUCN/SSC, 2013); they can be effective in preventing the extinction of species, re-establishing healthy populations (Bubac et al., 2019), and reversing rapid biodiversity loss (Seddon et al., 2007). In general, these translocations have mostly focused on mammals, birds, and reptiles (Bajomi et al., 2010; Bouma et al., 2020; Bubac et al., 2019; Fischer & Lindenmayer, 2000; Seddon et al., 2005); but the practice is not yet widely applied given difficulties like high costs and limited success rate in creating self-sustaining populations (Fischer & Lindenmayer, 2000; Griffith et al., 1989). Since successful translocations are more likely to be published, the success rate may be even lower than is evident from the literature (Miller et al., 2014).

Biodiversity loss continues despite widespread awareness of the problem and attempts to mitigate any threats (Butchart et al., 2010; Seddon et al., 2014). As a solution, the field of reintroduction biology (Moro et al., 2016; Seddon et al., 2007) has developed to examine and implement more integrative measures to maximise translocation success rates, including a broader inclusion of invertebrates to re-establish food webs. Insects vastly outnumber most other animal groups, suffer from habitat fragmentation and local extinction, and are probably more suitable for low-cost translocations than vertebrates (Hochkirch et al., 2007; Pearce Kelly et al., 1998). However, few insect conservation translocations have taken place compared to terrestrial vertebrate translocations (Bajomi et al., 2010). According to a recent study, 79 of 554 conservation translocations featured insects or other invertebrates (Bubac et al., 2019), of which most of them were focused on butterflies with a strong bias towards Europe and North America (Bellis et al., 2019; Nason et al., 2021). Additionally, few translocations have involved urban spaces (van Heezik & Seddon, 2018), even though invertebrates may be suitable for habitat restoration of these areas (e.g. Contos et al., 2021; Hannon & Hafernik, 2007).

Here, we examine the flightless ’Larapuna matchstick grasshopper’ (*Vandiemenella viatica*, Morabidae), a species <3 cm long with low mobility and dispersal capacity compared to winged insects: a characteristic that make them susceptible to local extinctions due to habitat reduction and modification (Weyer et al., 2012). This species is the only Morabidae species in Melbourne; it was formerly abundant but is now restricted to high-quality remnants of native vegetation. Conservation translocations to revegetated sites and other remnants in urban and suburban regions may be feasible for extending this species’ presence and assuring its survival in Melbourne. As far as we know, this is Australia’s first orthopteran translocation assessment. Most orthoptera translocations are from New Zealand (Bellis et al., 2019; Nason et al., 2021), where giant Wētā crickets have a long history of conservation management (Watts et al., 2008).

Long-term evaluations are typically considered more reliable when assessing the success of a reintroduction, as early performance may not be an accurate indicator of future viability (Seddon, 2013). However, there is no generally accepted approach for determining success (Robert et al., 2015; Seddon, 1999, 2015), and specific criteria for success will depend on the species being reintroduced and the context being considered as well as practical constraints. For instance, Jourdan et al. (2019) considered studies successful upon observing a new generation emerge after reproduction of the released individuals, while Bellis et al. (2019) defined a translocation as successful if the population released persisted at the release site beyond the species’ lifecycle duration, and if the latest monitoring reveals the persistence of the population at the release site. Our study also defined a successful translocation based on the detection of at least one individual after one generation, but we acknowledge that detection of our target species is imperfect, and that a full understanding of species establishment requires different phases of population development to be considered, including establishment, growth, and regulation (as discussed by Seddon (2013)).

Instead, this study focussed on early establishment and involved two different sets of experiments, a one-week trial of six releases and 36 translocations evaluated over a year. The primary objective of the one-week trial was to determine the detection probability of individuals based on various factors, such as effort, sex, colour, and marking effects. With the 36 translocations, we aimed to (1) investigate the short-term success of 36 urban translocations while providing preliminary data for future studies; (2) estimate the probability of detection and occupancy of *V. viatica* including site and survey covariates; (3) examine the individuals distance and direction travelled in a short period as background information for our monitoring design; and (4) examine the correlation of site covariates and number of released individuals that could influence the short-term translocation success in urban environments as assessed by the abundance and presence of individuals. Overall, the study aimed to provide insights into the early establishment of *V. viatica* in urban environments and those covariates that could influence this process.

## 2. Materials and methods

### 2.1. The species

*V. viatica* is currently found from the Eyre Peninsula (South Australia) to Bairnsdale (Victoria), from the coast to the interior of the mainland of about 400 km, and in certain northern and eastern Tasmanian coastal locations (Figure 1a). It is found in coastal heathy habitats (White et al., 1964) and open woodlands, spanning a broad diversity of plants. Although it is not monophagous, it prefers certain groups of plants (Blackith & Blackith, 1966). For instance, the species is found in open heathland with beard-heaths (*Leucopogon*) and Tassel Rope-rush (*Hypolaena*) in the Royal Botanic Gardens, Cranbourne, in the eastern part of the metropolitan Melbourne region (MMR); grassland with flax-lilies (*Dianella*) and everlasting daisies (*Chrysocephalum*) in Truganina Cemetery, in the west of the MMR; and open woodland with *Cassinia* and *Kunzea* shrubs and several grasses in Diamond Creek, in the northeast area of the MMR (Figure 1b).

**Figure 1.**
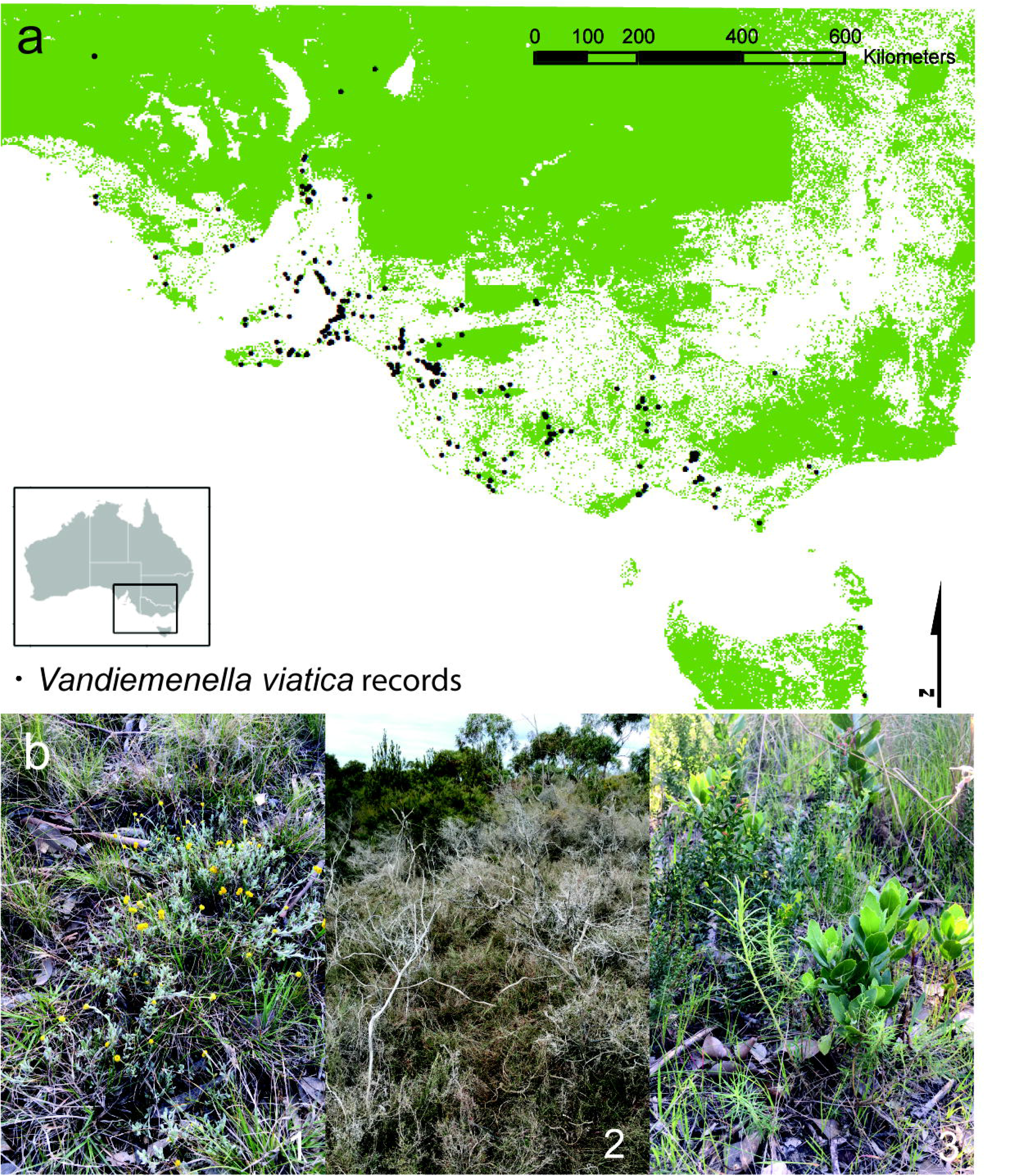
a. Extant Major Vegetation Groups (MGV) (present native vegetation) and *V. viatica* historical and current records (Data Source: MGV-National Vegetation Information System V6.0 © Australian Government Department of Agriculture, Water and the Environment. *V. viatica* records-Atlas of Living Australia occurrence download at https://doi.org/10.26197/ala.d80320c6-ad15-46a6-9e31-d672db19e26b. Accessed 26 May 2022). b. *V. viatica* natural habitat vegetation types in 1. Truganina Cemetery; 2. Royal Botanic Gardens Cranbourne and 3. Diamond Creek.

*V. viatica*, like other morabine grasshoppers, has several colour morphs (Key, 1976) that match their environment (Figure A1a), making them hard to see when they are not jumping or moving. As an annual winter species, it starts its life cycle as an egg during late spring and early summer (Blackith & Blackith, 1969), hatching at the beginning of the year (White et al., 1967). Males mature in May and June, but females need another month or two (Blackith & Blackith, 1969). Adults may be spotted until late November or early December. Like other morabines, *V. viatica* shows chromosomal variation across its distribution (Kawakami, 2008); based on male chromosomes, the most common races are *viatica*19 (19 chromosomes) and *viatica*17 (17 chromosomes) groups are considered races, not species, since their genitalia are identical (White et al., 1964). Greater Melbourne has only the *viatica*19 chromosomal race (White et al., 1964). Despite chromosomal type, this species’ mtDNA differs greatly among populations (Kawakami et al., 2007, 2011). Recent SNP-based data on the threatened and related morabine *Keyacris scurra* (Hoffmann et al., 2021) showed that populations from this group might have considerable genetic divergence, and *V. viatica* support this (Hoffmann et al., 2023).

Although *V. viatica* is not categorised as endangered, vegetation degradation and land clearing for development (including residential areas, industrial development, infrastructure projects, and sports fields) are reducing their numbers. Based on extrapolations from the species records (Atlas of Living Australia, 2022a) to the Major Vegetation Group model before and after 1750 (National Vegetation Information System V6.0, Department of Climate Change, Energy, the Environment and Water, 2020), we found that 52% of its natural habitat has now been lost throughout its distribution (Figure 1a). For the Melbourne area, the percentage lost is 93% (Figure A2). Moreover, dense weedy vegetation has invaded much of the remaining habitat, making it unsuitable for *V. viatica*. Several genetically distinct populations of the species have been identified, including one located in the Truganina Cemetery (Hoffmann et al., 2023). Despite the constant threat of urban development, this population persists within a small patch of native grassland.

### 2.2. Study area and release criteria selection

We used the Ecological Vegetation Classes (EVCs) (Victorian Department of Environment and Primary Industries, 2014) (Figure A2) and published information on *V. viatica* preferred habitat and food plants to identify sites around metropolitan Melbourne potentially suitable for translocations. We applied the following selection criteria: (1) *V. viatica* absence as determined from surveys by 3 experienced collectors in 30 minutes under suitable weather conditions (air temperatures ≥12 °C), (2) presence of plant species that could serve as shelter (native grasses) and food (daisies and/or other herbs and shrubs), (3) open grasslands or woodlands with heath and with partial sun exposure (canopy cover ≤ 80%), (4) absence of dense weed cover, and (5) accessibility.

We found 36 suitable translocation sites, consisting of three remnants and 33 revegetated areas between 47 m^2^ to 12 000 m^2^ in size (Figure 2). The revegetated areas consisted of native grasses and herbs recently established in public parks next to footpaths and recreational grassy areas. The remnants would have been heavily impacted by human activities in the past but had not been completely invaded by weeds and still supported native grasses and daisies.

**Figure 2.**
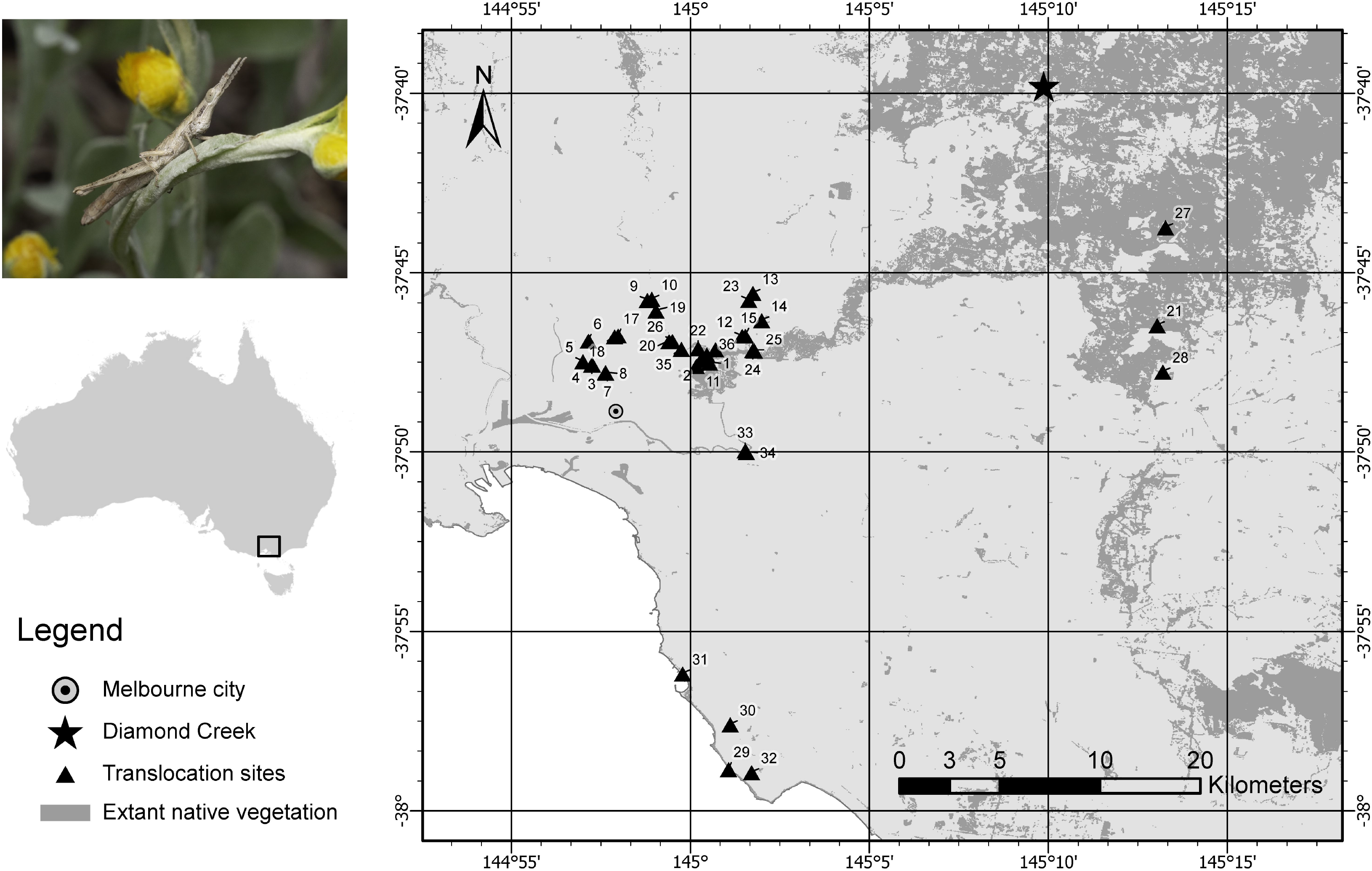
Release sites for *V. viatica* sourced from the Diamond Creek population. Extant native vegetation was taken from Ecological Vegetation Classes: Copyright ©The State of Victoria, Department of Environment, Land, Water & Planning. Numbers correspond to site names in Table A1.

### 2.3. Source populations, collection, manipulation, and release of individuals

The source population was collected from a privately owned property of approximate 8.7 hectares in Diamond Creek, corresponding to the EVC of Grassy Dry Forest (EVC 22). Because of its unsuitability for agriculture, this EVC has persisted across much of this area, including some extensive remnants (Bull & Sinclair, 2014). The land was slated for home building and had a population density of approximately 1-2 grasshoppers per m^2^ in suitable habitat patches. We visited the site five times and used a pooter to collect an average of 403 individuals each day (total of 2014) between August and September 2021. A total of 1851 individuals were transferred to portable cages of 20 x 20 x 20 cm and separated by colour and sex in the lab to then kept in groups of mostly 50 for translocation, consisting of the same number of females and males and a similar colour proportion as found in the field. Only three individuals from the 1851 died before release. Overall, the release period covered a wetter than-average period with temperatures close to average for Greater Melbourne (Bureau of Meteorology, 2022).

Within three days of being collected, we released 25 individuals in each of two courtyard sites, 50 to 72 individuals in each of 33 larger sites, and 142 individuals in one of the larger sites (Table A1). After collection, individuals were provided water and food plants from the source site (mostly *Cassinia*). The releases were carried out on days when temperatures were sufficient for grasshopper activity (>10°C) to allow them to locate refuges and avoid predators.

The one-week trial consisted of six releases in a managed revegetation area with daisies and native grasses along the Yarra River at Burnley, an inner-city suburb (next to site 33 and 34 in Figure 2). In total, 300 individuals, 150 males and 150 females, were released (50 per site); 150 of these were marked with acrylic paint using a six-dot identification code (Walker & Wineriter, 1981) to test the feasibility of this technique for marking *V. vandiemenella* (Figure A1b).

### 2.4. Surveys

After taping off a 10-meter radius circle around the release site, the first, second, and third surveys were conducted. The area was plotted through a satellite image for reference and signposted in the field. Searching took 45 minutes until the entire area was covered. One person could thoroughly search a 1 m^2^ area in approximately 15 seconds. An additional 15 minutes was spent searching for adequate vegetation within 20 m radius of the release site. Releases in linear and smaller areas used a 10 m radius circular area, but the searching time was then decreased to less than 45 minutes (see Figure 3). The search radii were based on a dispersal study where the furthest distance travelled by an individual was 11.28 m in 12 days (Blackith & Blackith, 1969). For the final survey, we expanded the search area to cover to surrounding habitat with suitable vegetation. Limits were set by cement footpaths, tracks wider than 3 m, or fences. To eliminate observer bias, sites with no grasshoppers were revisited by the same surveyor and by a second surveyor if the grasshopper was still undetected.

**Figure 3.**
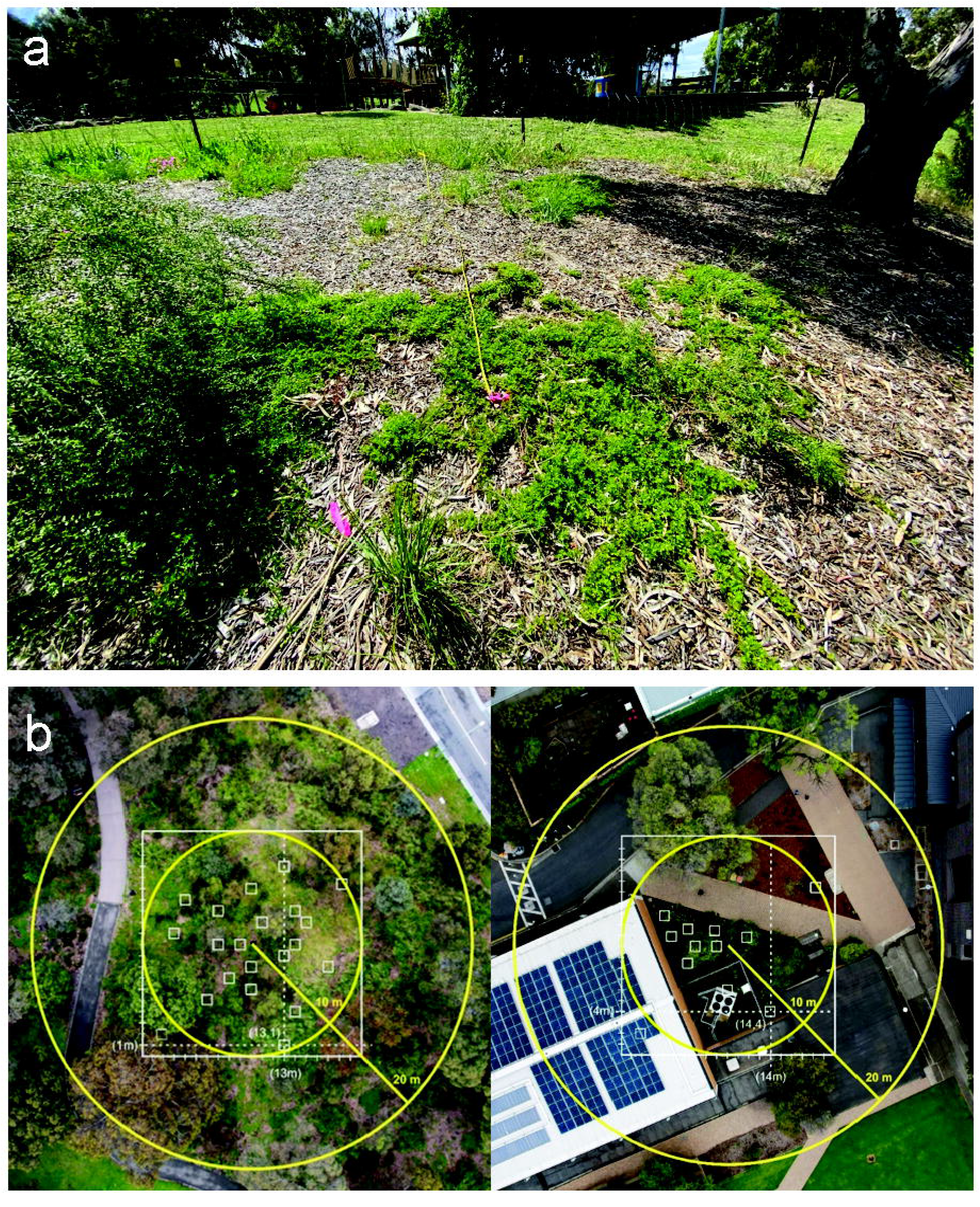
Survey design. a. 10 m radius from release point. b. survey areas and quadrants for vegetation cover analysis.

### 2.5. Site and survey covariates

The habitat site was characterised by components likely to affect individual establishment and detection. These included habitat area, canopy cover (CC), understorey life forms cover (ULFC), plant species richness, weed cover, and known food cover (Table 1). Satellite images (Google Earth Pro v7.3.4) were used to compute the habitat area that corresponded to adequate vegetation delimited by the barriers noted above. CC was considered because it influences numerous biological processes affected by light, water availability, wind, and temperature (Lemmon, 1956). We assessed ULFC using the “Habitat hectares scoring methods” (DSE, 2004), which refers to plant groups with similar three-dimensional structure and comparable overall dimensions (DSE, 2004). To determine ULFC and species cover, 20 random 1 m x 1 m quadrants were established inside a 10 m circular region from the release point. The quadrant was positioned using two random numbers smaller than 20 to represent horizontal and vertical axes (Figure 3b). The measurements were made between December – January 2022 when no adults were present. The vegetation composition results are summarized in Table A2. For analysis, the number of released individuals was also considered as a site covariate.

**Table 1.** List of variables considered to characterise habitats.

| Variable | Abbreviation | Units | Possible effect on individuals |
| --- | --- | --- | --- |
| Habitat area |  | m<sup>2</sup> | Dispersal |
| Canopy cover | CC | 1-100 % | Life cycle |
| Understory life form cover | ULFC | 1-100 % |  |
| Shrub cover | Shrub |  | Food/Shelter |
| Herb cover | Herb |  | Food/Shelter |
| Graminoid cover | Gram |  | Shelter |
| Moss cover | M |  | Food |
| Litter cover | L |  | Shelter/Oviposition |
| Bare soil cover | BS |  | Oviposition |
| Bark mulch cover | BM |  | Shelter/Oviposition |
| Rocks and logs cover | RL |  | Shelter/Oviposition |
| Plant species richness | PSR |  | Food/Shelter |
| Weed cover | Weed | 1-100 % | Life cycle |
| Known food cover | Known food | 1-100 % | Food |

The survey covariates were soil temperature (C°) at 0 cm of depth, and survey season. The soil temperature data was obtained from the microclimate model of NicheMapR (Kearney & Porter, 2017) using the ERA5 reanalysis dataset (Hersbach et al., 2020) for the 2021-2022 period. The model was validated against data provided by the Bureau of Meteorology, Melbourne Airport station. To determine the timing of our four surveys, we considered overall species generation time and projected development stages for each season. The first survey was conducted one month after release (November) and targeted the original parental generation (F0). The second survey was carried out in late summer (February and March) when one-month-old F1 nymphs were expected and were large enough to be detected. The third survey was conducted in late autumn around the beginning of June, targeting grasshoppers at the third/fourth instar. Finally, exactly one year after the release, we conducted our last survey (August and September), which targeted F1 adults (Table A1). Below these surveys are referred to by season.

During the surveys, rainfall levels were higher than average, except for the summer survey in February, a month with less than 25% of the average rainfall across Greater Melbourne (Bureau of Meteorology, 2022). Mean maximum temperatures in 2022 were close to the annual averages for the region (Bureau of Meteorology, 2022).

### 2.6. Statistical analysis

Analyses were performed in R 4.2.1 (R Core Team, 2022) using the Rstudio environment (RStudio Team, 2022) and plots were made using the R package ’ggplot2’ (Wickham, 2014), except where indicated.

For the short-term trials, we obtained an individual’s detection probability three days after release with a one person-hour sampling effort and one week later with a 10 person-hours sampling effort. A two-way ANOVA was performed to analyse the effect of release sites, effort, sex, colour, and marking on detection probability. For the 36 site translocations, we conducted occupancy analysis to estimate the probability of detection for *V. viatica* using the package ‘unmarked’ (v1.2.5 Fiske & Chandler, 2011). Occupancy analysis is a widely used method for estimating the occurrence and detection of species in ecological studies (MacKenzie et al., 2002). Given the short-term nature of our study, we fitted a single-species, single-season occupancy model to the data. To account for potential sources of variation in the probability of detection, we incorporated site and survey covariates into the model. We assessed the model using goodness-of-fit tests and model selection criteria. We used the package MuMIn version 1.46.0 to rank and select a model based on the Akaike Information Criteria (AIC) in terms of AICc (corrected for small sample sizes), delta AIC (ΔAIC), and its weight (w) (Barton, 2022). Kullback-Leibler distance is used by the AIC to select an optimal model (Burnham & Anderson, 2002). Only models with delta AICs equal to or below two were considered of comparable quality (Symonds & Moussalli, 2011).

In our study of the 36 translocation sites, we used a Friedman test (run in IBM Statistics ver. 28) assess differences among sites and surveys in the number of grasshoppers collected. Additionally, we aimed to determine whether the first survey’s results were related to those of the final survey. To do this, we computed a non-parametric Kendall’s rank correlation coefficient comparing the probability of detecting F0 individuals with the number of grasshoppers in the final survey (F1). We also employed Fisher’s exact test to compare grasshopper detection rates in the first and final surveys. Finally, we fitted a negative binomial model (estimated using maximum likelihood (ML)) to predict the number of detected individuals in the last survey (Spring – 2022) as a function of the number of released individuals. We excluded sites where individuals were never found based on the assumption that unsuitable habitat conditions, rather than a lack of individuals released, were responsible for the absence of grasshoppers across all surveys at a particular site.

We additionally hypothesised that more grasshoppers would be detected at higher temperatures and sunny conditions. To test this, we computed a negative binomial regression using the function ‘glm.nb()’ of the package MASS (Ripley et al., 2022), relating soil temperature (C°) at 0 cm of depth to the number of individuals detected for each survey and a combination of all surveys. We also included additional secondary surveys in these analyses but excluded sites where grasshoppers were never recovered. Note that data from the short term trials were not included in these analyses given that conditions were similar across the one-week period considered.

In each survey, we measured the distance and direction of an individual’s position relative to the release point. Since the data violated normality assumptions even after transformation, Kruskal-Wallis H and Mann–Whitney U tests were used to compare distances between sites and between males and females. Comparisons were made for both the short-term trials and for data obtained after one year in the case of the translocations. A Rayleigh test of uniformity was used to analyse movement direction, and a Mardia-Watson-Wheeler test was used to compare females and males; for these analyses, we used the R package ‘circular’ (Lund et al., 2022). Polar graphs were made using the R package ‘plotly’ (Sievert et al., 2022).

Finally, we used Generalised Linear Models (GLMs), including negative binomial regression and multiple logistic regression, to examine the association of abundance and presence/absence data respectively, with site covariates and number of released individuals. For these analyses, all covariates were tested for normality using Shapiro-Wilk statistics and standardised for parameter comparisons using the z-score (Quinn & Keough, 2002). We reduced covariates due to the small number of data points (Neter et al., 1996) (n=36). First, we used Spearman’s correlations (Elledge et al., 2013) and a correlation matrix to test for collinearity and remove duplicate covariates, but none were found. Covariates were also analysed using principal components analysis (PCA) (e.g. Kelt et al., 1994), but principal components were not used because they did not explain >50% of the total variance. Finally, we used a forward regression selection of covariates in each season using the ’regsubsets’ function of the R package ’leaps’, limiting the number of predictors to six (Gram, PSR, BM, Known food, Herb, and Weed) for abundance and seven (CC, Shrub, Gram, M, BS, Weed, and PSR) for presence/absence. We later ranked and selected models based on ΔAIC as mentioned before for occupancy and detection probability.

## 3. Results

### 3.1 Abundance and presence of grasshoppers

Our four resurveys yielded 20, 26, 18 and 26 positive sites with grasshoppers, and 68, 56, 39, and 99 individuals, respectively (Table A1). When increasing the size of our survey area in the final survey, we found two more positive sites (n=28) and 14 extra individuals (n=113), meaning that 77.78% of sites were positive one year after translocation. Grasshoppers were never found at six of the sites. The number of grasshoppers varied significantly among surveys (Friedman test, *X*^2^ = 79.63, df = 4, *p* < 0.001) and among sites (*X^2^* = 99.34, df = 35, *p* < 0.001), with the highest numbers detected in the last survey (Spring-2022) at the end of August and start of September when F1s were surveyed (Figure 4).

**Figure 4.**
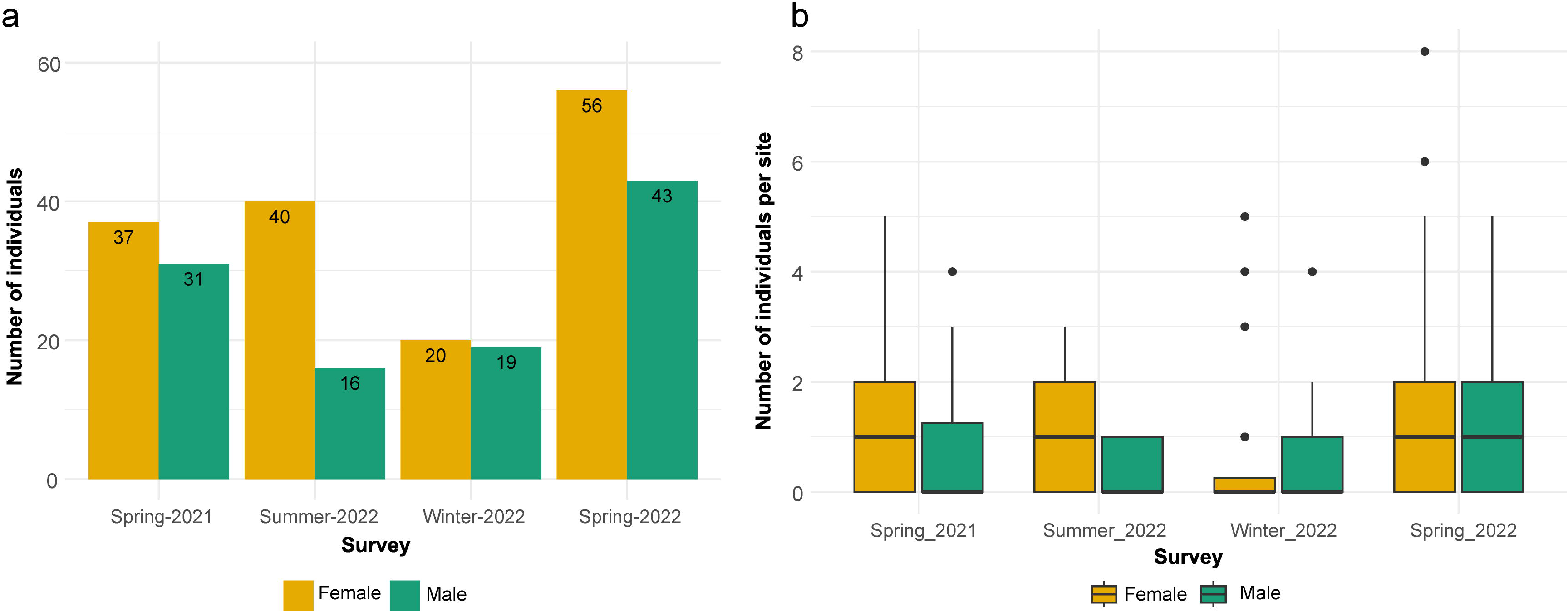
a. Total number of females and males detected in each survey. b. Boxplot of number of females and males detected per site by survey season.

### 3.2 Occupancy, detection, and movement

For the short-term trial, a two-way ANOVA revealed that effort had a statistically significant effect on the mean detection probability (*F*_1,5_ = 18.03, *p* = 0.008), but there was no effect of release site (*F*_5,5_ = 2.056, *p* = 0.224). Sex did not have a significant effect on the mean detection probability (*F*_1, 17_ =3.144, *p* = 0.09), and neither did the release site (*F*_5,17_ = 1.133, *p* = 0.381), with no interaction (*F*_5,12_ =0.245, *p* = 0.935). Colour did not affect the mean detection probability (*F*_1,17_ = 0.408, *p*= 0.531), and neither did release site (*F*_5,17_ = 1.602, *p* = 0.213), and there was no interaction between site and colour (*F*_5,12_ = 0.269, *p* = 0.922). Also, marking did not have a statistically significant effect on the detection probability (*F*_1,17_ = 0.114, *p* = 0.74) nor release site (*F*_5,17_ = 1.079, *p* = 0.407).

For the 36 translocations, we found an occupancy probability of 0.83 (SE = 0.06, 95% CI [0.68, 0.92]) and a detection probability of 0.74 (SE = 0.04, 95% CI [0.65, 0.81]) without considering survey variability. In other words, our study predicted *V. viatica* to occur in about 83% of the sites, and to detect the species, when present, about 74% of the time. When considering survey occasions, the estimated probability of detection of the last survey was the highest (*p* = 0.93, SE = 0.05, CI [0.76, 0.98]), followed by the second survey (*p* = 0.83, SE = 0.07, CI [0.65, 0.93]), the first survey (*p* = 0.67, SE = 0.09, CI [0.48, 0.81]) and finally the third survey (*p* = 0.53, SE = 0.09, CI [0.36, 0.7]). The results of model selection for *V. viatica* are shown in Table 2. The best model included the covariates graminoid cover, plant species richness, and weed cover to explain the probability of occupancy, while soil temperature was informative on the probability of detection (Figure A3a).

**Table 2.** Models of occupancy and detection probability for *V. viatica* using the Akaike information criterion (AICc), Delta AIC and Akaike weight (w). Only models with ΔAICc < 2 are shown.

| Model | df | AICc | Delta | w |
| --- | --- | --- | --- | --- |
| $\psi$ (Gram, PSR, Weed) p(temp) | 6 | 162.2 | 0.00 | 0.37 |
| $\psi$ (Gram, PSR, Weed) p(.) | 5 | 163.5 | 1.29 | 0.19 |
| $\psi$ (Gram, PSR) p(temp) | 5 | 164.0 | 1.81 | 0.15 |

We found that females had a similar occupancy probability (*ψ =* 0.82 SE=0.07, 95% CI [0.65, 0.92]) to males (*ψ* = 0.77, SE= 0.07, 95% CI [0.42, 0.89]). Pooling across surveys, the probability of detection of females (*p* = 0.63, SE = 0.05, 95% CI [0.53, 0.71]) was similar to that of males (*p* = 0.59, SE = 0.05, 95% CI [0.49, 0.69]). When including survey times as a factor, the trend was similar for females and males, with higher values in the last survey for females (*p* = 0.78, SE = 0.08, 95% CI [0.6, 0.9]) and males (*p* = 0.8, SE = 0.08, 95% CI [0.6, 0.91]), and smaller values in the third survey for females (*p* = 0.31, SE = 0.09, 95% CI [0.17, 0.49]) and males (*p* = 0.44, SE = 0.1, 95% CI [0.27, 0.63]).

Kendall’s rank correlation analyses revealed a weak positive but non-significant correlation between the F0 individuals’ detection probability in the first survey and the number of F1 individuals detected in the last survey (τ = 0.21, *p* = 0.12). One year after translocation, we detected F1 individuals in all the 20 positive sites noted in the first survey, detected eight more positive sites, and found that six sites negative in the first survey were also negative in the last survey. A Fisher’s exact test on these numbers was highly significant (*p* = 0.002), rejecting the null hypothesis that detection in the two surveys was independent. The number of detected individuals in the last survey (Spring – 2022) was unrelated to the number of released individuals (negative-binomial model, *R^2^* = 0.05, β = 0.16, 95% CI [-0.12, 0.51], *p* = 0.28). However, it should be noted that similar numbers were released at most sites.

For the one-week trial, there were significant differences in dispersal distance among sites within a week (Kruskal-Wallis H test, *X^2^*_5_ *=* 32.493; *p* < 0.001) but males and females did not differ in the distance moved (Mann–Whitney U test, *W*=5872.5, *p* = 0.583). After one year, females and males did not differ in mean distances from the release site at our 36 translocation locations (*W* = 1487.5, *p* = 0.967). The distances found are shown in Table 3. In one of the sites, two F1 individuals, one female and one male were found more than 40 m from the release site, requiring F1 or F0 individuals to cross a three-meter gravel path. The maximum distance travelled by marked individuals was 11.11 m, with a minimum of 0.6 m after one week. One individual returned to the release point after travelling 2 m, while others returned close to it, suggesting an exploratory behaviour. There was a significant positive linear relationship between normalised habitat area and maximum dispersal distance from one generation to the other (*r* = 0.47, N = 28, p = 0.009) (Figure 5).

**Table 3.** Distance from release site (m)

|  | Minimum | Maximum | Mean |
| --- | --- | --- | --- |
| *One-week after release* |  |  |  |
| Female | 0.1 | 11 | 1.81 m (SD = 1.98) |
| Male | 0 | 9.45 | 1.67 m (SD = 1.64) |
| *One-year after release* |  |  |  |
| Female | 0.8 | 49.7 | 11.19 m (SD=11.34) |
| Male | 0.2 | 55.9 | 10.85 m (SD=11.29) |

**Figure 5.**
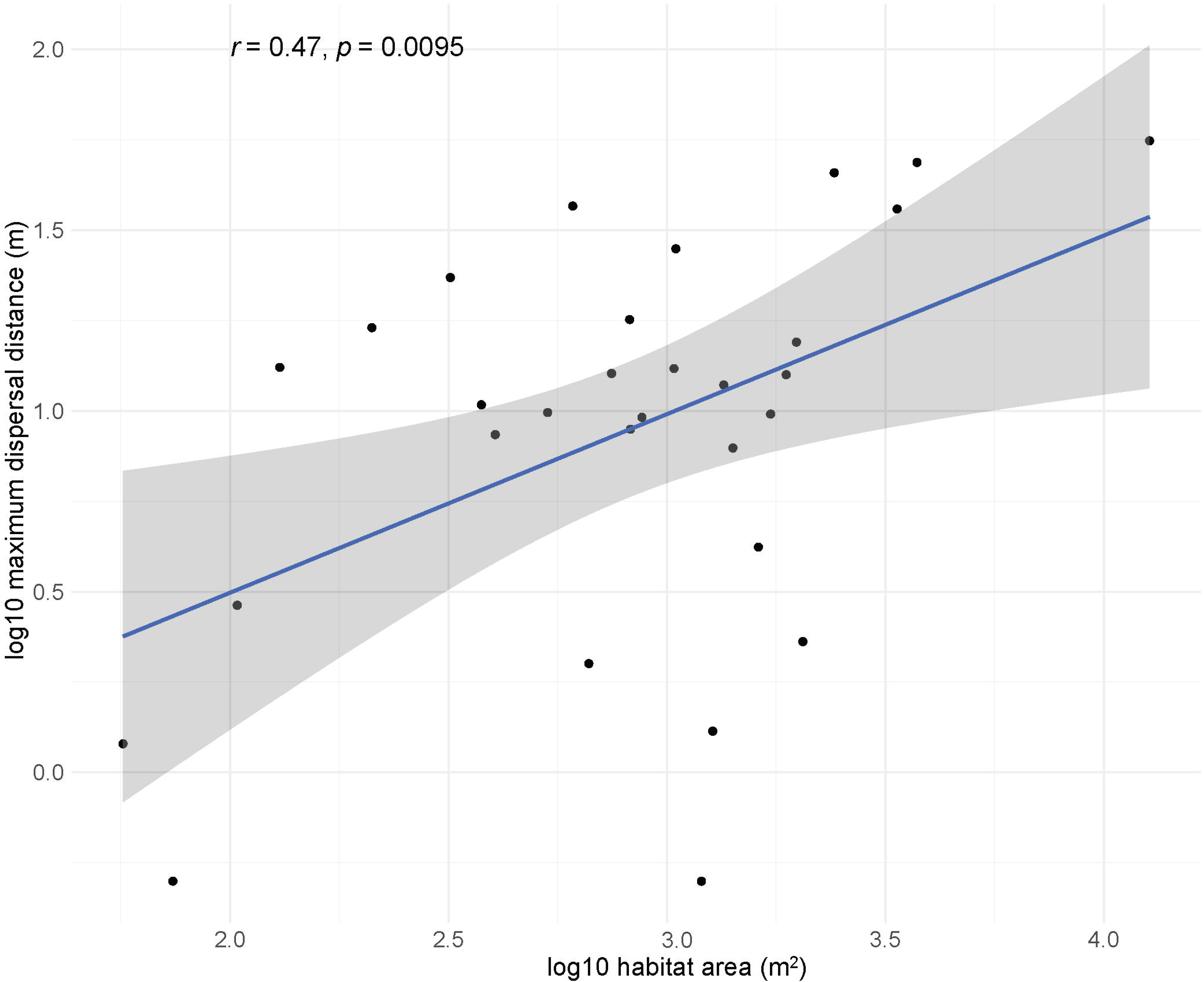
Relationship between habitat area and maximum dispersal distance across the 36 translocation sites (log_10_ scale).

The Rayleigh test showed that in the short-term trial, the grasshoppers’ movement direction was not uniform (θ = 156.996°, Rayleigh test: r = 0.1568, *p*=0.005, n = 215), but it was uniform between generations for the 36-site translocation (θ = 188.932°, Rayleigh test: r = 0.1191, *p* = 0.207, n = 111). The Mardia–Watson-Wheeler test for homogeneity of angles showed no significant difference between the direction of movements of females and males for the short-term trial (*W* = 1.1204, df = 2, *p* = 0.571) and the 36-site translocations (*W* = 0.42148, df = 2, *p* = 0.81) (Figure 6).

**Figure 6.**
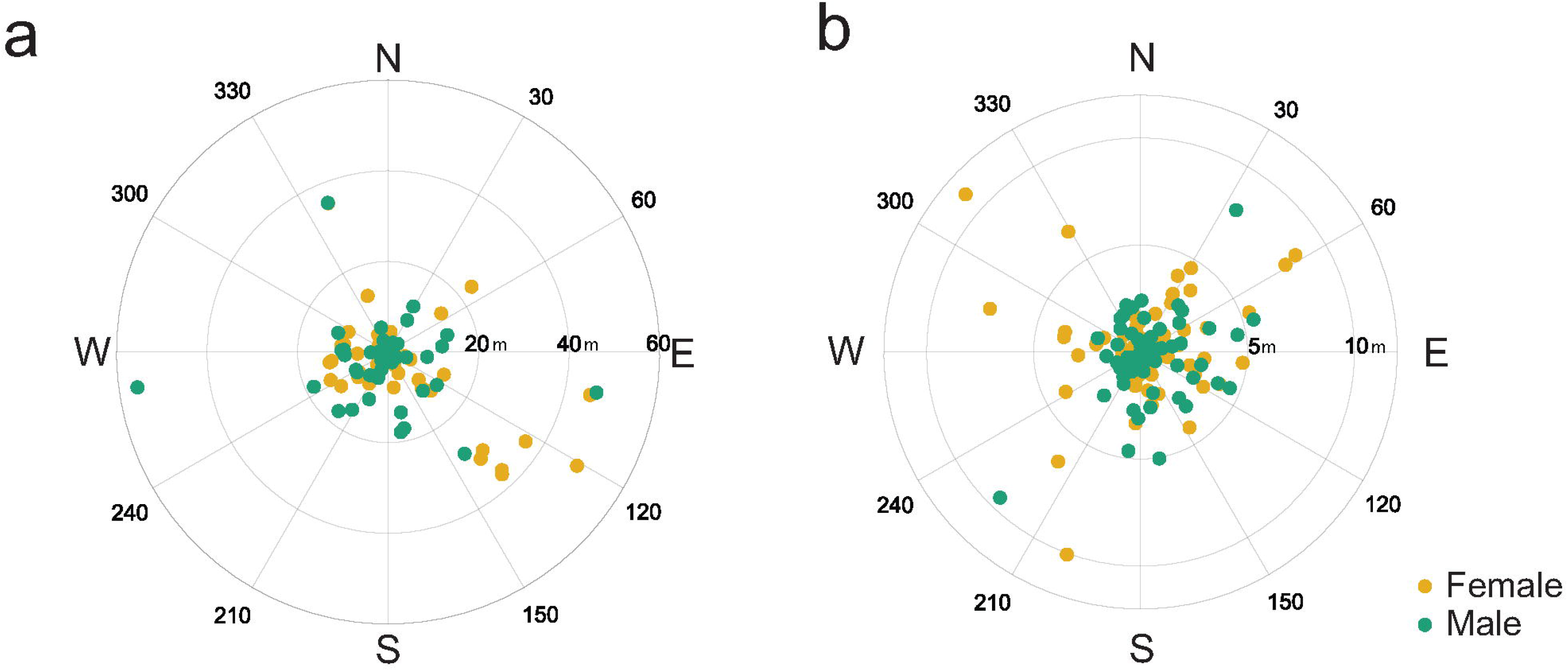
Scatter plot in polar coordinates showing the distance and direction travelled by females and males. a. in the 36 translocation sites after a year and b. in the Burnley release after a week.

Only the third survey (Winter – 2022) showed a significant effect of estimated soil temperature on the number of individuals detected (R^2^_N_ = 0.34, β = 0.68, 95% CI [0.22, 1.19], *p* = 0.002) (Figure 7).

**Figure 7.**
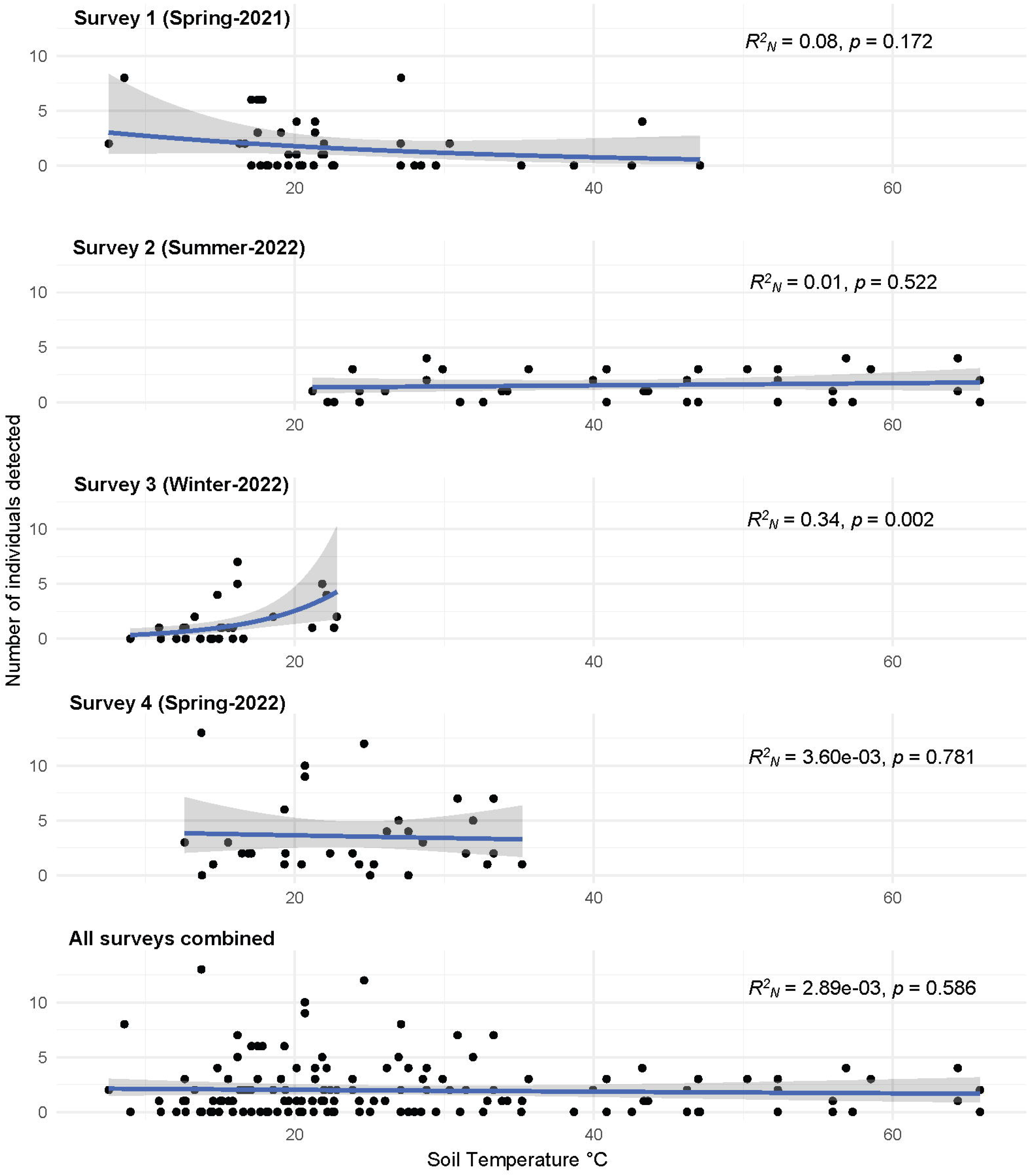
Relationship between the number of individuals detected and estimated soil surface temperature in each survey fitted with a negative-binomial regression model.

### 3.3 Habitat models and model selection

Sixty-four models for number of individuals and 128 for *V. viatica* presence were obtained from all explanatory variable combinations. Considering individual abundance, we obtained substantial explanatory power models for the four surveys (R^2^_N_ = 0.78, R^2^_N_ = 0.59, R^2^_N_ = 0.48, and R^2^ = 0.55). Graminoid cover was statistically significant for the first (β = -0.74, 95% CI [-1.27, -0.25], *p* = 0.003), third (β = -0.71, 95% CI [-1.40, -0.09], *p* = 0.028), and fourth (β = -0.44, 95% CI [-0.88, -0.02], *p* = 0.039) survey; weed cover was statistically significant for the first (β = -1.28, 95% CI [-2.58, -0.36], *p* =0.027) and last surveys (β = - 0.82, 95% CI [-1.71, -0.20], *p* =0.031); and plant species richness was statistically significant for the first (β = 0.53, 95% CI [0.16, 0.93], *p* = 0.004), and second surveys (β = 0.41, 95% CI [0.12, 0.71], *p* = 0.006). Considering the individuals presence, we obtained substantial explanatory power for the four surveys (R^2^ = 0.64, R^2^ = 0.49, R^2^ = 0.29, R^2^ = 0.55), where graminoid cover effect was statistically significant for the first survey (β = -0.94, 95% CI [-1.81, -0.34], *p* = 0.01), while moss cover (β = -1.60, 95% CI [-3.67, -0.27], *p* = 0.046) and plant species richness (β = 3.35, 95% CI [1.02, 7.25], *p* = 0.026) were statistically significant for the last survey. Simple regressions of abundance and presence/absence against the variables with the highest Akaike weight are displayed in Figure A3. The AICc, ΔAICc < 2 and w outcomes considering all the explanatory variables possible combinations are shown in Table 4. Considering the abundance and presence of grasshoppers in every survey, we found that the explanatory variables common among surveys were graminoid cover and the plant species richness.

**Table 4.** Model selection table explaining the abundance and presence of grasshoppers using the Akaike’s information criterion (AICc), Delta AIC and Akaike weight (w) in the different survey seasons. Only models with ΔAICc < 2 are shown.

| | AICc | $\Delta AIC$ | w |
| --- | --- | --- | --- |
| **Spring 2021** |  |  |  |

|  | AICc | ΔAIC | w |
| --- | --- | --- | --- |
| Abundance of grasshoppers |  |  |  |
| Gram + PSR + Weed | 121.8 | 0.00 | 0.208 |
| Gram + Herb + PSR + Weed | 122.2 | 0.42 | 0.169 |
| Gram + BM + PSR + Weed | 122.5 | 0.78 | 0.141 |
| Gram + BM + Herb + PSR + Weed | 123.3 | 1.50 | 0.098 |
| Presence of grasshoppers |  |  |  |
| Gram + M + PSR | 33.5 | 0.00 | 0.114 |
| Gram + PSR | 33.6 | 0.07 | 0.110 |
| Gram + M + PSR + Weed | 33.9 | 0.37 | 0.095 |
| Gram + PSR + Weed | 35.0 | 1.48 | 0.054 |
| **Summer 2022** |  |  |  |
| Abundance of grasshoppers |  |  |  |
| Herb + PSR + Weed | 107.1 | 0.00 | 0.180 |
| Known food + PSR + Weed | 107.5 | 0.43 | 0.145 |
| PSR + Weed | 108.6 | 1.50 | 0.085 |
| Presence of grasshoppers |  |  |  |
| M + Shrub + Weed | 37 | 0.00 | 0.063 |
| PSR + Shrub + Weed | 37.1 | 0.10 | 0.060 |
| M + PSR + Shrub + Weed | 37.3 | 0.32 | 0.053 |
| PSR + Weed | 37.4 | 0.44 | 0.050 |
| **Winter 2022** |  |  |  |
| Abundance grasshoppers |  |  |  |
| Gram + PSR | 102.0 | 0.00 | 0.143 |
| Gram | 102.9 | 0.86 | 0.093 |
| Gram + Herb + PSR | 103.6 | 1.57 | 0.065 |
| Gram + Herb | 103.8 | 1.82 | 0.058 |
| Presence of grasshoppers |  |  |  |
| PSR + Shrub | 48.1 | 0.00 | 0.066 |
| CC + Gram | 49.1 | 0.97 | 0.041 |
| CC + Shrub | 49.2 | 1.08 | 0.039 |
| Graminoid | 49.4 | 1.31 | 0.035 |
| **Spring 2022** |  |  |  |
| Abundance of grasshoppers |  |  |  |
| Gram + Herb + Weed | 153.2 | 0.00 | 0.155 |
| Herb + Weed | 154.2 | 1.06 | 0.091 |
| Gram + Weed | 154.3 | 1.14 | 0.088 |
| Presence of grasshoppers |  |  |  |
| Moss + PSR + Shrub | 35.8 | 0.00 | 0.081 |
| Gram + PSR | 36.0 | 0.17 | 0.075 |
| Gram + M + PSR + Shrub | 36.1 | 0.28 | 0.071 |
| Gram + PSR + Shrub | 36.7 | 0.91 | 0.052 |

## 4. Discussion

Our research suggests insect translocations as a promising conservation tool in urban landscapes, supporting the notion that conservation initiatives in urban Australia might assist rare and vulnerable endemic insect species (New, 2018). Despite limited ecological knowledge of *V. viatica* and no translocation precedent for this grasshopper, we achieved a short-term translocation success of 77.78% after one year, covering an entire life cycle of the species. Although longer-term post-release monitoring is fundamental to assessing translocation success (IUCN/SSC, 2013), translocations are more likely to fail in the early stages and short-term evaluations are useful in providing preliminary insights into the effectiveness of the reintroduction as noted by Bubac et al. (2019). We were also able to test how species attributes (sex and colour) and environmental conditions (soil temperature) impact their detection, how much *V. viatica* spreads following release, and how habitat characteristics affect grasshopper occupancy, abundance and presence.

### Detection and movement

Grasshopper detection may depend on the size of individuals and sexual dimorphism. For instance, Schori et al. (2020b) found that the detectability of the grasshopper *Brachaspis robustus* was correlated with body length and sex. Females of that species are 2-3 times larger than males and are more likely to be found than males. In the case of *V. viatica*, early male and female nymphs are around the same size with dimorphism developing in the 3^rd^-4^th^ instar, with females taking an extra moult to become about one centimetre larger than males. Despite this, we did not find sex effects in the detection probability later in the season, indicating that size may not be a critical issue in its detection.

Activity in ectotherms is susceptible to environmental temperature at the microclimate scale (Bramer et al., 2018; Kearney et al., 2021). Two different studies by Schori et al. (2020a; 2020b) found that ground temperatures >13.6°C did not affect the detection probability of the grasshopper *B. robustus*, although cloud cover did have a significant effect in one of their studies (Schori et al., 2020a). Our study showed that *V. viatica* were detected under air temperature range from 10.9 to 30.7 °C and soil temperature of 7.6 to 65.8 °C. As a winter species, we might expect *V. viatica* detectability to be affected by cold weather, consistent with the significant and positive effect we found only during the winter surveys . These grasshoppers are highly cryptic and are mainly detected when they hop. They appear to be more likely to hop on the ground than when they are perched or nestled in vegetation, a behaviour that requires them to be warm enough to be active before descending to the ground. Thus substrate temperature (which incorporates air temperature, wind speed, and solar radiation (Kearney & Porter, 2017)) should be recorded to monitor the species, and is easily measured with an IR thermometer in the field. Additionally, biophysical models for predicting field body temperatures in different microhabitats could be used to help design monitoring systems in the future (e.g., Saleeba et al. (2020)).

Our study suggests expanding the sample region beyond the 20 m radius we first employed. Depending on patch area and habitat availability, individuals travelled about 25 m (over two months) from their release location. In Sites 1 and 20 (see Table A1), where the initial food plants dried up, F1 individuals were found more than 40 m from the release site in patches of daisies and beauty-heads, suggesting movement in response to resource availability. Half of the release locations were close to a concrete cyclist/pedestrian path or a two-lane road, explaining the non-uniformity in the movement direction. We found no sex differences in movement rates, even though males may go further to increase mating chances (Hochkirch et al., 2007). A lack of sex difference is consistent with previous observations on this species (Blackith and Blackith, 1969) and other flightless grasshoppers (Weyer et al. 2012) and consistent with our finding that sexes were equally detectable. Finally, we found no directionality in movement, suggesting that these grasshoppers track suitable habitats.

Individuals may move ambidirectionally when encountering unfavourable environments (Kindvall, 1999), which occurred in our short-term trial when some marked individuals returned to food plants at the original release point.

### 4.1. Habitat suitability

Sufficient food is a crucial component of habitat quality for translocations (McGrath et al., 2017). We discovered that grasshopper occupancy, abundance and presence increased with plant species richness, perhaps because grasshoppers had more feeding options in case one plant species senesced. Different *V. viatica* life stages may favour different plant species. Graminoid cover had a negative effect on the occupancy, abundance and presence of grasshoppers; at Sites 2 and 13, grasses grew while other plant species withered. Also, while grasses and weeds offer cover, a high density will limit grasshopper thermoregulation. *V. viatica* may thrive in areas with rich plant diversity and short grasses with cover <50%.

Understanding life stages is essential for translocations (Samways, 1994). We exclusively released adults because females deposited eggs shortly after release, maximising reproductive output. A possible disadvantage of this strategy is that it may decrease late-stage grasshoppers’ ability to learn about local food and shelter resources. The effectiveness of our translocations over one year suggests that grasshoppers avoided high levels of predation and parasitism, located good oviposition sites, and acquired enough food to develop successfully to adulthood in the ensuing generation. In some cases, grasshoppers reached suitable food plants patches tens of metres away from the release point, perhaps through olfaction (Hopkins & Young, 1990). Despite our short-term success, it would be worth exploring the merits of releasing hatchlings or early instars.

### 4.2. Future concerns

Long-term population survival is still unclear and supplementation through more releases may be needed. Random mortality prior to the establishment of a sustainable population may prevent efficient colonisation (Sa Berggren, 2001). If mortality decreases population size, the resulting small populations may be susceptible to extinction due to genetic factors (Hoffmann et al., 2021; Lynch et al., 1995). One possible approach that could be explored involves utilizing a number of individuals from diverse subpopulations which may prevent the loss of genetic diversity and lower the risk of inbreeding as demonstrated in *Gryllus campestris* by Witzenberger & Hochkirch (2008). *V. viatica*’s minimal viable population size for long-term persistence requires additional investigation to inform future introductions. Also, more work is needed to assess mortality caused by local predators like spiders (which prey on young grasshoppers) and birds/lizards/mantids (which can prey on adults) (Wagner et al., 2005).

Source populations and their suitability for certain translocation sites need more investigation. Our source population was from a habitat with a high diversity of plant species (1284 plant species, historical and recent records – 5 km radius Atlas of Living Australia, 2022b). Does that make them more suitable for translocations than those individuals found in more restricted habitats, like populations at Truganina cemetery (589 plant species, historical and recent records – 5 km radius Atlas of Living Australia, 2022c)? Does the genetic variability within the species make some populations more adaptable and suitable for reintroduction purposes?

The only viable long-term solution to habitat loss is the implementation of active conservation and management practices (Schultz & Hammond, 2003). Revegetation initiatives must incorporate insect and food plant phenology to ensure grasshopper survival. As an example, translocation Site 2 only had one potential food plant, the annual herb *Helichrysum luteoalbum -* Jersey Cudweed. It disappeared during summer, leaving individuals without any possible food resources. In contrast, at sites with the perennial *Chrysocephalum*, *V. viatica* had available food during its whole life cycle. Because graminoids and weeds were negatively correlated with the grasshoppers’ number and presence, active management through weeding may be necessary at grassy release sites.

Despite some challenges, our findings showed that insect translocations are viable, even in small urban spaces of revegetated areas, with a fraction of the time and resources required for vertebrate translocation initiatives that need substantial investment over extended periods (Gedir et al., 2004). For example, a translocation project for the black-footed rock-wallaby (*Petrogale lateralis*) cost AU$3.86 million over 11 years (Ireland et al., 2018); Hilbers et al. (2020) predicted a cost of ∼US$3 million to reintroduce 1753 captive-reared peregrine falcons (*Falco peregrinus*) in California to restore and maintain a minimum viable population; a project reintroducing the green and golden bell frog (*Litoria aurea*) tadpoles had an approximate overall cost of AU$190K for more than four years (Daly et al., 2008). In contrast, our translocations cost < AU$1K in non-labour costs, with around AUS$20K for surveying, collecting and monitoring. Insects are excellent candidates for translocation based on their life-history characteristics (Bellis et al., 2019), comparatively straightforward logistics of catching, transporting, and releasing individuals, and their potential ability for fast population growth once established in a new place (New, 2012). Also, their small size allows them to be released in smaller areas than vertebrates, reducing pre- and post-release habitat management costs (Bellis et al., 2019).

## 5. Conclusion

Invertebrates have been historically overlooked in translocation research for conservation purposes, despite their immense potential, particularly beyond reintroductions in the wild. Our work on morabine grasshoppers shows that urban insect translocations are possible, and that short-term monitoring is crucial. Based on our findings, we recommend survey designs to augment detection, that includes life-history characteristics and habitat requirement of the species that affect the success of establishing the released individuals. The benefits of this process include the re-establishment of a unique species that was once common in the area, flow-on effects to other faunal biodiversity given that the morabine grasshoppers are a seasonally important source of food for other animals, and an opportunity to inform the public about insect conservation around cities.

## CRediT authorship contribution statement

Hiromi Yagui: Conceptualization, Methodology, Investigation, Data curation, Formal analysis, Writing – original draft, Writing – review & editing. Michael R. Kearney: Conceptualisation, Funding acquisition, Methodology, Investigation, Project administration, Writing – review & editing, Supervision. Ary A. Hoffman: Conceptualisation, Funding acquisition, Methodology, Investigation, Project administration, Writing – review & editing, Supervision.

## Declaration of competing interest

The authors declare no conflict of interest that could influence the work presented in the manuscript.

## Data availability

Open access through melbourne.figshare.com

DOI: 10.26188/22009346

The DOI becomes active when the item is published.

## Acknowledgements

HY was funded by Melbourne Research Scholarship from the University of Melbourne. MRK and AAH received funding from the Australian Research Council, Discovery Grant DP190100990, the Victorian Department of Land, Water and Planning, and the Melbourne City Council. We are grateful to A. Liu, S. Jaboor, K. Kaufmann, and A. Tello for their fieldwork support, and also to S. Guo, E. Pearce, J. Sadler, N. Wenhrynowycz, S. Falkenberg, N. Phuong Nguyen, N. Nguyen, D. Noonan-O’Keeffe, C. Stanley, and A. Loebmann, participants in the Burnley one-week trial. We acknowledge the valuable feedback and suggestions provided by the reviewers and associate editor, which have greatly contributed to the improvement of our work.

## Notes

### Competing Interest Statement

The authors have declared no competing interest.

