## Supplementary_information for "Restoring declining species through translocations: A test case using flightless grasshoppers in an urban setting"

**Appendix:**

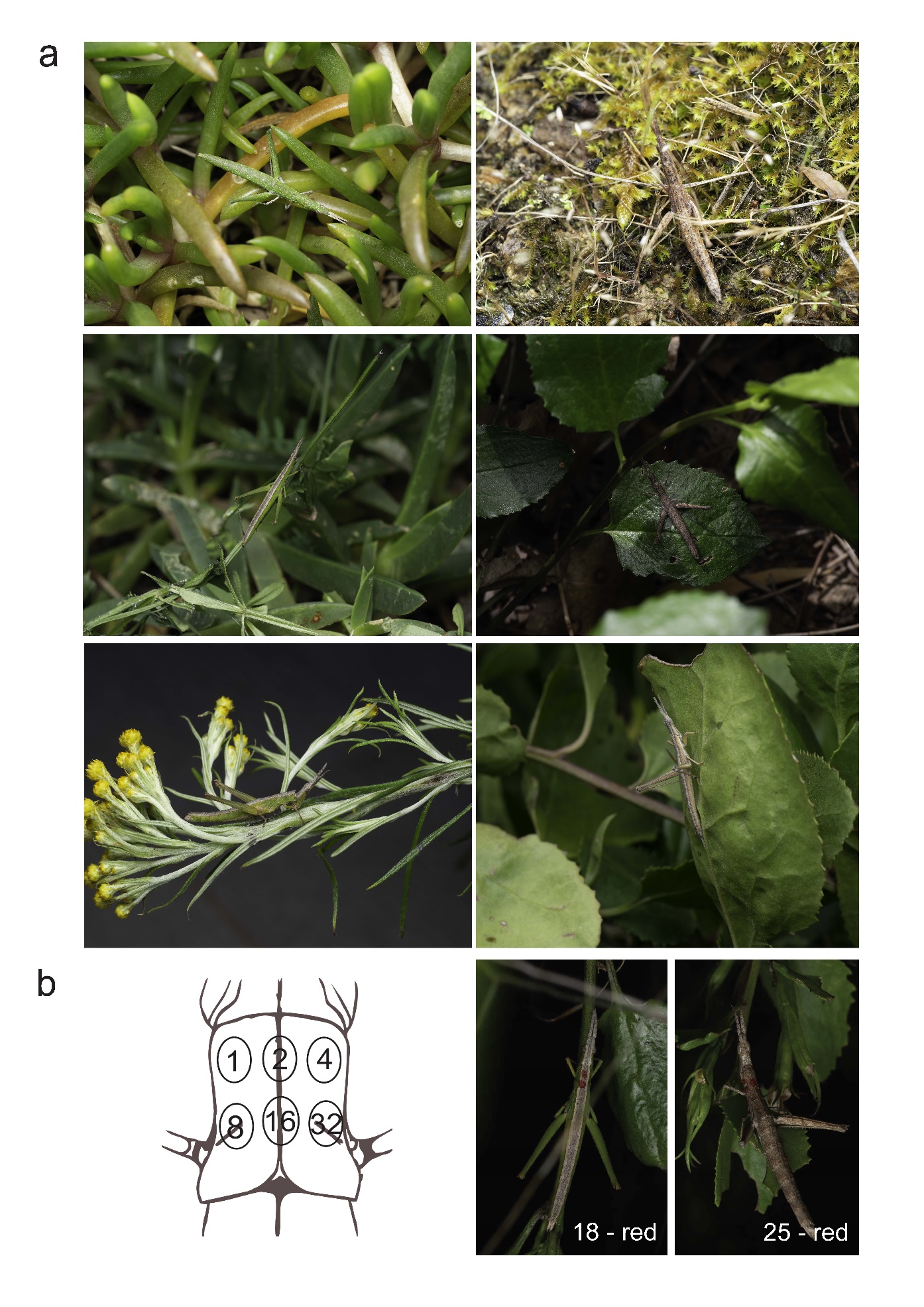

**Figure A1** a. Examples of *Vandiemenella viatica* colour morphs. b. Marking design based on Walker & Wineriter, 1981

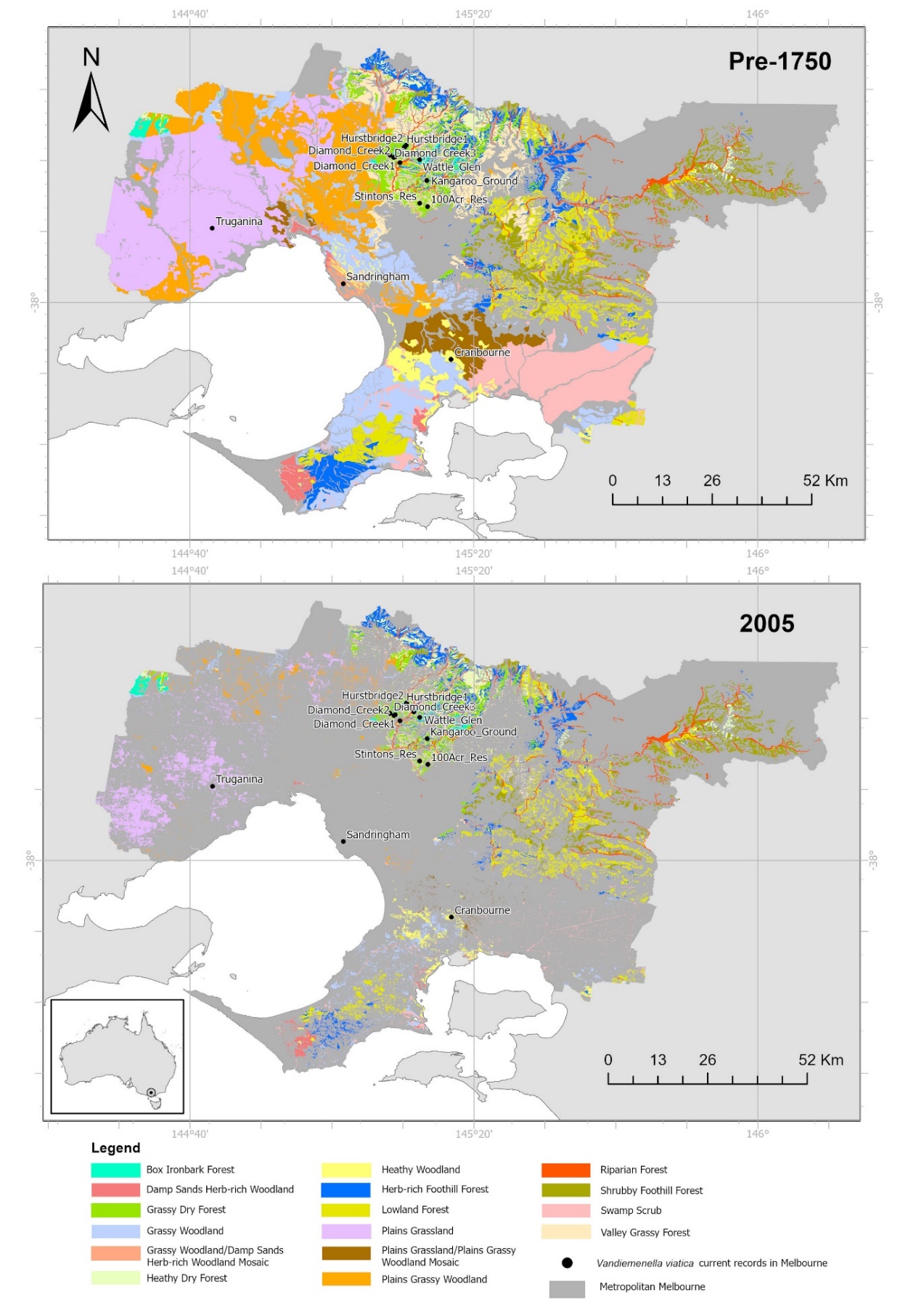

**Figure A2** Current *Vandiemenella viatica* population records in the Melbourne region and their corresponding pre-1750 and 2005 native vegetation classified into Major Vegetation Groups in Metropolitan Melbourne (Copyright ©The State of Victoria, Department of Environment, Land, Water & Planning).

**Table A1** Number of individuals collected in each translocations site according to season and type

| **N** | **Translocation site** | **Category** | **Bioregion** | **Area** | **Number of released individuals** | **Spring_2021** | **Summer_2022** | **Winter_2022** | **Spring_2022** | **Spring_2022 (Beyond 20m radius)** | **Mean** | **SE** |
| --- | --- | --- | --- | --- | --- | --- | --- | --- | --- | --- | --- | --- |
| 1 | Westfield South Oval | Revegetated | Victoria Volcanic Plain | 12738 | 142 | 0 | 1 | 1 | 4 | 7 | 2.6 | 1.29 |
| 2 | Clifton Hill | Revegetated | Victoria Volcanic Plain | 2027.05 | 50 | 0 | 0 | 0 | 0 | 0 | 0.0 | 0.00 |
| 3 | Royal Park (Bayles Street Grassland) -1 | Revegetated | Gippsland Plain | 763.98 | 50 | 0 | 0 | 0 | 0 | 0 | 0.0 | 0.00 |
| 4 | Royal Park (Bayles Street Grassland) -2 | Revegetated | Gippsland Plain | 876.96 | 50 | 2 | 3 | 5 | 5 | 5 | 4.0 | 0.63 |
| 5 | Royal Park (Close to tram stop) | Revegetated | Gippsland Plain | 319.6 | 50 | 8 | 4 | 7 | 11 | 13 | 8.6 | 1.57 |
| 6 | Royal Park (Close to train station) | Revegetated | Gippsland Plain | 1038.51 | 50 | 0 | 0 | 0 | 2 | 2 | 0.8 | 0.49 |
| 7 | Unimelb (next to Gym) | Courtyard flowerbed | Gippsland Plain | 103.91 | 25 | 0 | 0 | 1 | 4 | 4 | 1.8 | 0.92 |
| 8 | Unimelb (Infront of Gym) | Courtyard flowerbed | Gippsland Plain | 74.01 | 25 | 0 | 1 | 0 | 0 | 0 | 0.2 | 0.20 |
| 9 | Jones Park, Brunswick East | Revegetated | Victoria Volcanic Plain | 376.23 | 40 | 0 | 3 | 1 | 2 | 2 | 1.6 | 0.51 |
| 10 | Velodrome, Brunswick East | Revegetated | Victoria Volcanic Plain | 69.15 | 50 | 0 | 0 | 0 | 0 | 0 | 0.0 | 0.00 |
| 11 | Field street | Revegetated | Victoria Volcanic Plain | 1200.34 | 72 | 7 | 3 | 0 | 2 | 2 | 2.8 | 1.16 |
| 12 | Wingrove west | Revegetated | Victoria Volcanic Plain | 56.91 | 50 | 0 | 1 | 0 | 1 | 1 | 0.6 | 0.24 |
| 13 | Mansfield | Revegetated | Victoria Volcanic Plain | 662.84 | 50 | 0 | 1 | 0 | 0 | 0 | 0.2 | 0.20 |
| 14 | Darebin parklands | Revegetated | Victoria Volcanic Plain | 1273.7 | 50 | 0 | 1 | 0 | 1 | 1 | 0.6 | 0.24 |
| 15 | Wingrove east | Courtyard flowerbed | Victoria Volcanic Plain | 56.88 | 50 | 0 | 0 | 0 | 0 | 0 | 0.0 | 0.00 |
| 16 | Fairfield boathouse | Revegetated | Victoria Volcanic Plain | 1728.26 | 50 | 0 | 2 | 1 | 2 | 2 | 1.4 | 0.40 |
| 17 | Hardy Gallagher Reserve_Park_Street | Revegetated | Victoria Volcanic Plain | 1350.18 | 50 | 1 | 1 | 0 | 2 | 2 | 1.2 | 0.37 |
| 18 | Hardy Gallagher Reserve_Lang_Street | Revegetated | Victoria Volcanic Plain | 533.55 | 50 | 2 | 1 | 0 | 3 | 3 | 1.8 | 0.58 |
| 19 | Merri Creek Blyth Street, Brunswick East | Revegetated | Victoria Volcanic Plain | 47.37 | 50 | 0 | 0 | 0 | 0 | 0 | 0.0 | 0.00 |
| 20 | Inner Circle-Thomas Kidney Reserve | Revegetated | Victoria Volcanic Plain | 3737.59 | 50 | 4 | 2 | 0 | 0 | 2 | 1.6 | 0.75 |
| 21 | Domeney Reserve | Revegetated | Highlands-Southern Fall | 3366.2 | 50 | 1 | 3 | 0 | 0 | 3 | 1.4 | 0.68 |
| 22 | Hall Reserve (North) Playground | Revegetated | Victoria Volcanic Plain | 2416.78 | 50 | 4 | 3 | 0 | 1 | 2 | 2.0 | 0.71 |
| 23 | Clarendon | Revegetated | Victoria Volcanic Plain | 404.77 | 50 | 2 | 3 | 1 | 12 | 12 | 6.0 | 2.47 |
| 24 | Alphington Park I | Revegetated | Victoria Volcanic Plain | 1620.28 | 48 | 6 | 2 | 1 | 1 | 1 | 2.2 | 0.97 |
| 25 | Alphington Park II | Revegetated | Victoria Volcanic Plain | 1416.11 | 49 | 0 | 1 | 1 | 1 | 1 | 0.8 | 0.20 |
| 26 | Merri Creek Linear Reserve | Revegetated | Victoria Volcanic Plain | 609.11 | 50 | 1 | 0 | 0 | 1 | 3 | 1.0 | 0.55 |
| 27 | Blooms Rd. | Remnant | Highlands-Southern Fall | 130.02 | 50 | 4 | 3 | 1 | 2 | 2 | 2.4 | 0.51 |
| 28 | Loughies Bushland Reserve | Revegetated | Highlands-Southern Fall | 1979.95 | 50 | 3 | 2 | 4 | 4 | 4 | 3.4 | 0.40 |
| 29 | Beach Road | Revegetated | Gippsland Plain | 2047.9 | 50 | 1 | 0 | 2 | 1 | 1 | 1.0 | 0.32 |
| 30 | Royal Avenue | Revegetated | Gippsland Plain | 825.41 | 50 | 3 | 0 | 1 | 1 | 1 | 1.2 | 0.49 |
| 31 | Hampton Beach | Revegetated | Gippsland Plain | 541.23 | 50 | 0 | 0 | 0 | 0 | 0 | 0.0 | 0.00 |
| 32 | Donald Macdonald Reserve | Remnant | Gippsland Plain | 747.52 | 50 | 2 | 1 | 1 | 5 | 5 | 2.8 | 0.92 |
| 33 | Burnley_Site1 | Revegetated | Gippsland Plain | 821.94 | 50 | 3 | 4 | 5 | 9 | 9 | 6.0 | 1.26 |
| 34 | Burnley_Site6 | Revegetated | Gippsland Plain | 211.25 | 50 | 6 | 4 | 4 | 10 | 10 | 6.8 | 1.36 |
| 35 | Merri Creek_close to bridge | Revegetated | Victoria Volcanic Plain | 1876.42 | 50 | 6 | 3 | 0 | 6 | 6 | 4.2 | 1.20 |
| 36 | Merri Creek Trail_reveg | Revegetated | Victoria Volcanic Plain | 1048.76 | 50 | 2 | 3 | 2 | 6 | 7 | 4.0 | 1.05 |

**Table A2** List of the 132 plant species recorded, their life form, weediness, and number of translocation sites where they are found. Weediness was categorised according to the EVC bioregion benchmark © The State of Victoria Department of Sustainability and Environment (2004), available at <https://www.environment.vic.gov.au/biodiversity/bioregions-and-evc-benchmarks>

| **Species code** | **Life form** | **Weediness** | **Number of sites** |
| --- | --- | --- | --- |
| *Acacia_paradoxa* | Medium Shrub | - | 2 |
| *Acacia_sp1* | Medium Shrub | - | 5 |
| *Acacia_sp2* | Medium Shrub | - | 1 |
| *Acaena_echinata* | Medium Herb | - | 2 |
| *Acaena_novae_zelandiae* | Medium Herb | - | 6 |
| *Acrotriche* | Prostrate Shrub | - | 1 |
| *Allium* | Medium to Tiny Non-tufted Graminoid | Weed | 1 |
| *Alternanthera* | Medium Herb | - | 2 |
| *Arthropodium* | Large Herb | - | 5 |
| *Asparagus asparagoides* | Scrambler or Climber | Weed | 1 |
| *Atriplex* | Medium Herb | - | 1 |
| *Austrostipa* | Large Tufted Graminoid | - | 3 |
| *Banksia* | Medium shrub | - | 1 |
| *Bolboschoenus* | Medium to Tiny Non-tufted Graminoid | - | 1 |
| *Bossiaea* | Medium Shrub | - | 2 |
| *Brachyscome_sp1* | Medium Herb | - | 6 |
| *Brachyscome_sp2* | Medium Herb | - | 1 |
| *Brachyscome_sp3* | Medium Herb | - | 1 |
| *Briza_maxima* | Medium to Small Tufted Graminoid | Weed | 3 |
| *Bromus* | Medium to Small Tufted Graminoid | Weed | 2 |
| *Brunonia_australis* | Small or Prostrate Herb | - | 2 |
| *Bulbine* | Medium to Small Tufted Graminoid | - | 1 |
| *Calocephalus* | Medium Shrub | - | 2 |
| *Carpobrotus* | Small or Prostrate Herb | - | 5 |
| *Cassinia_sp1* | Medium Shrub | - | 4 |
| *Cassinia_sp2* | Medium Shrub | - | 1 |
| *Cerastium* | Medium Herb | Weed | 1 |
| *Chrysocephalum_apiculatum* | Medium Herb | - | 13 |
| *Chrysocephalum_semipapposum* | Large Herb | - | 17 |
| *Coronidium* | Medium Herb | - | 1 |
| *Correa* | Medium Shrub | - | 2 |
| *Cotula* | Small or Prostrate Herb | - | 1 |
| *Crassula* | Small or Prostrate Herb | - | 1 |
| *Delairea* | Scrambler or Climber | Weed | 1 |
| *Dianella* | Medium to Small Tufted Graminoid | - | 18 |
| *Dichelachne* | Large Tufted Graminoid | - | 1 |
| *Dichondra* | Small or Prostrate Herb | - | 9 |
| *Disphyma* | Small or Prostrate Herb | - | 1 |
| *Drosanthemum* | Medium Herb | - | 1 |
| *Drosera* | Small or Prostrate Herb | - | 1 |
| *Echium_candicans* | Medium Shrub | - | 1 |
| *Einadia* | Medium Herb | - | 12 |
| *Enchylaena* | Small Shrub | - | 10 |
| *Epilobium* | Large Herb | - | 1 |
| *Eryngium* | Large Herb | - | 2 |
| *Eucalyptus* | Medium shrub | - | 4 |
| *Euphorbia_rigida* | Medium shrub | - | 1 |
| *Festuca_glauca* | Medium to Small Tufted Graminoid | - | 1 |
| *Ficinia* | Large Tufted Graminoid | - | 5 |
| *Frankenia* | Small Shrub | - | 1 |
| *Fumaria* | Medium Herb | Weed | 4 |
| *Galium* | Scrambler or Climber | Weed | 3 |
| *Gazania_rigens* | Medium Herb | - | 1 |
| *Genista* | Medium shrub | Weed | 1 |
| *Geranium* | Medium Herb | - | 8 |
| *Goodenia ovata* | Medium shrub | - | 4 |
| *Helichrysum_luteoalbum* | Medium Herb | - | 9 |
| *Hypericum* | Medium Herb | - | 2 |
| *Hypochoeris* | Medium Herb | Weed | 8 |
| *Juncus* | Large Tufted Graminoid | - | 2 |
| *Kunzea* | Medium shrub | - | 1 |
| *Lagurus_ovatus* | Medium to Tiny Non-tufted Graminoid | - | 1 |
| *Lasiopetalum* | Medium shrub | - | 1 |
| *Lepidium* | Large Herb | Weed | 1 |
| *Lepidosperma* | Medium to Small Tufted Graminoid | - | 1 |
| *Leptospermum* | Medium shrub | - | 5 |
| *Leucochrysum_albicans* | Medium Herb | - | 2 |
| *Leucophyta* | Small Shrub | - | 1 |
| *Linum* | Small or Prostrate Herb | Weed | 1 |
| LNG1 | Large Non-Tufted Graminoid | - | 2 |
| LNG2 | Large Non-Tufted Graminoid | - | 2 |
| LNG3 | Large Non-Tufted Graminoid | - | 2 |
| Lomandra | Large Tufted Graminoid | - | 11 |
| LTG1 | Large Tufted Graminoid | - | 2 |
| LTG2 | Large Tufted Graminoid | - | 1 |
| LTG3 | Large Tufted Graminoid | - | 1 |
| MH1 | Medium Herb | - | 1 |
| MH2 | Medium Herb | - | 1 |
| MH3 | Medium Herb | - | 1 |
| MH4 | Medium Herb | - | 2 |
| MH5 | Medium Herb | - | 1 |
| MNG1 | Medium to Tiny Non-tufted Graminoid | - | 1 |
| MNG2 | Medium to Tiny Non-tufted Graminoid | - | 2 |
| MNG3 | Medium to Tiny Non-tufted Graminoid | - | 11 |
| MNG4 | Medium to Tiny Non-tufted Graminoid | - | 5 |
| MNG5 | Medium to Tiny Non-tufted Graminoid | - | 8 |
| MNG6 | Medium to Tiny Non-tufted Graminoid | - | 1 |
| MTG1 | Medium to Small Tufted Graminoid | - | 1 |
| MTG2 | Medium to Small Tufted Graminoid | - | 1 |
| MTG3 | Medium to Small Tufted Graminoid | - | 4 |
| MTG4 | Medium to Small Tufted Graminoid | - | 2 |
| MTG5 | Medium to Small Tufted Graminoid | - | 2 |
| MTG6 | Medium to Small Tufted Graminoid | - | 4 |
| MTG7 | Medium to Small Tufted Graminoid | - | 2 |
| MTG8 | Medium to Small Tufted Graminoid | - | 1 |
| MTG9 | Medium to Small Tufted Graminoid | - | 3 |
| MTG10 | Medium to Small Tufted Graminoid | - | 1 |
| MTG11 | Medium to Small Tufted Graminoid | - | 1 |
| MTG12 | Medium to Small Tufted Graminoid | - | 1 |
| MTG13 | Medium to Small Tufted Graminoid | - | 1 |
| Myoporum | Medium shrub | - | 2 |
| Olearia | Medium shrub | - | 2 |
| Opercularia | Small or Prostrate Herb | - | 1 |
| Oxalis | Small or Prostrate Herb | - | 13 |
| Pelargonium | Medium Herb | - | 7 |
| *Pimelea* | Small Shrub | - | 1 |
| *Plantago* | Medium Herb | Weed | 9 |
| *Poa* | Medium to Small Tufted Graminoid | - | 10 |
| *Poranthera_microphylla* | Medium Herb | - | 3 |
| *Pycnosorus* | Large Herb | - | 2 |
| *Rhagodia* | Medium shrub | - | 4 |
| *Rubus* | Small Shrub | - | 2 |
| *Rytidosperma* | Medium to Small Tufted Graminoid | - | 15 |
| *Senecio* | Large Herb | - | 9 |
| SH1 | Small or prostrate herb | - | 2 |
| SH2 | Small or prostrate herb | - | 1 |
| SH3 | Small or prostrate herb | - | 1 |
| SH4 | Small or prostrate herb | - | 2 |
| *Sonchus* | Large Herb | Weed | 13 |
| *Stellaria* | Medium Herb | - | 1 |
| *Taraxacum* | Medium Herb | Weed | 1 |
| *Tetragonia* | Scrambler or Climber | - | 2 |
| *Themeda_triandra* | Medium to Small Tufted Graminoid | - | 10 |
| *Tragopogon* | Large Herb | - | 1 |
| *Trifolium_repens* | Small or prostrate herb | Weed | 4 |
| *Trifolium* | Small or prostrate herb | Weed | 3 |
| TTG1 | Tiny Tufted Graminoid | - | 3 |
| *Veronica* | Large Herb | - | 1 |
| *Vicia* | Small or prostrate herb | Weed | 2 |
| *Vittadinia* | Small or prostrate herb | - | 1 |
| *Wahlenbergia* | Medium Herb | - | 16 |
| *Xerochrysum* | Large Herb | - | 3 |

a
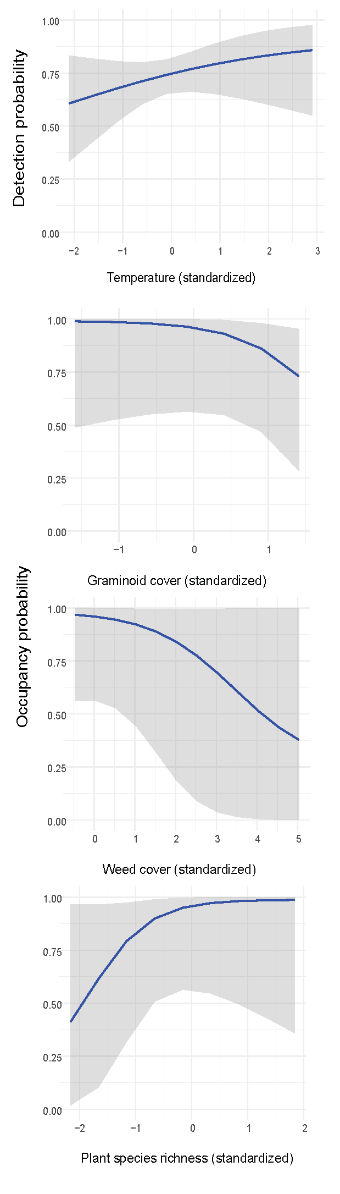
b
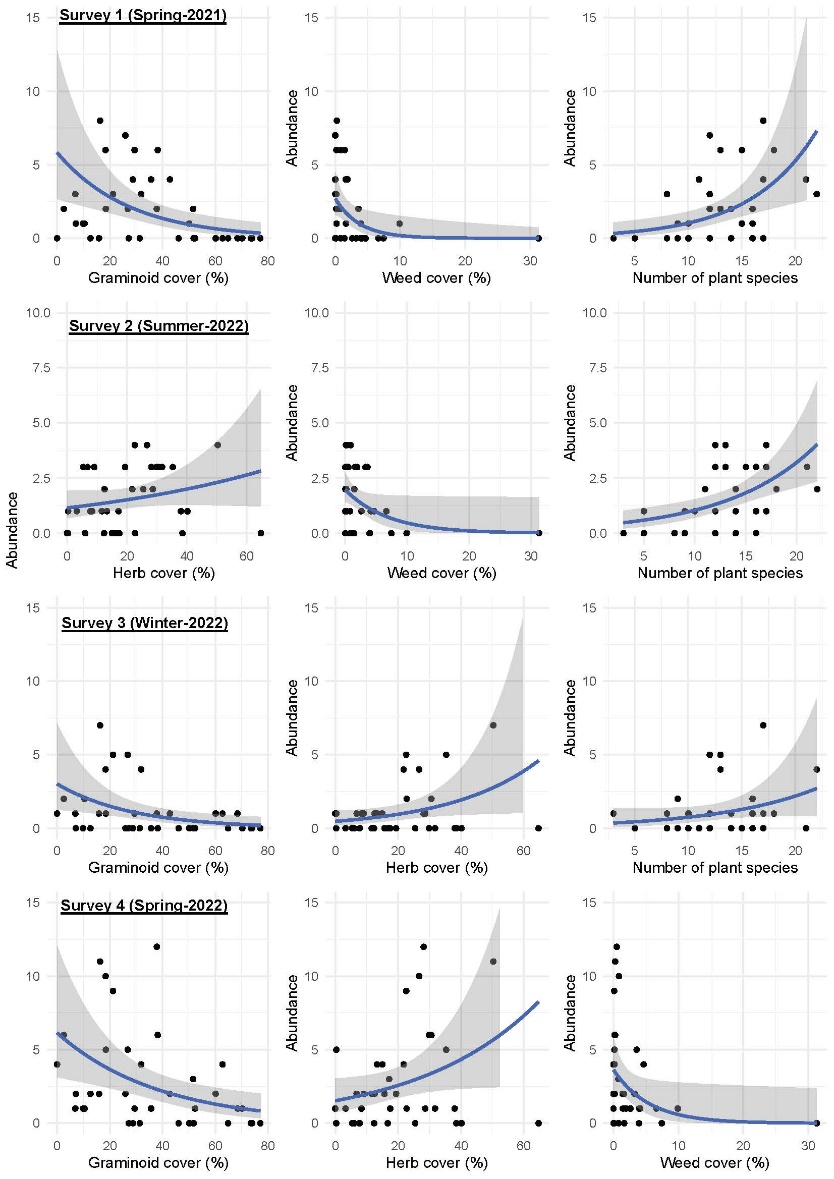
 c
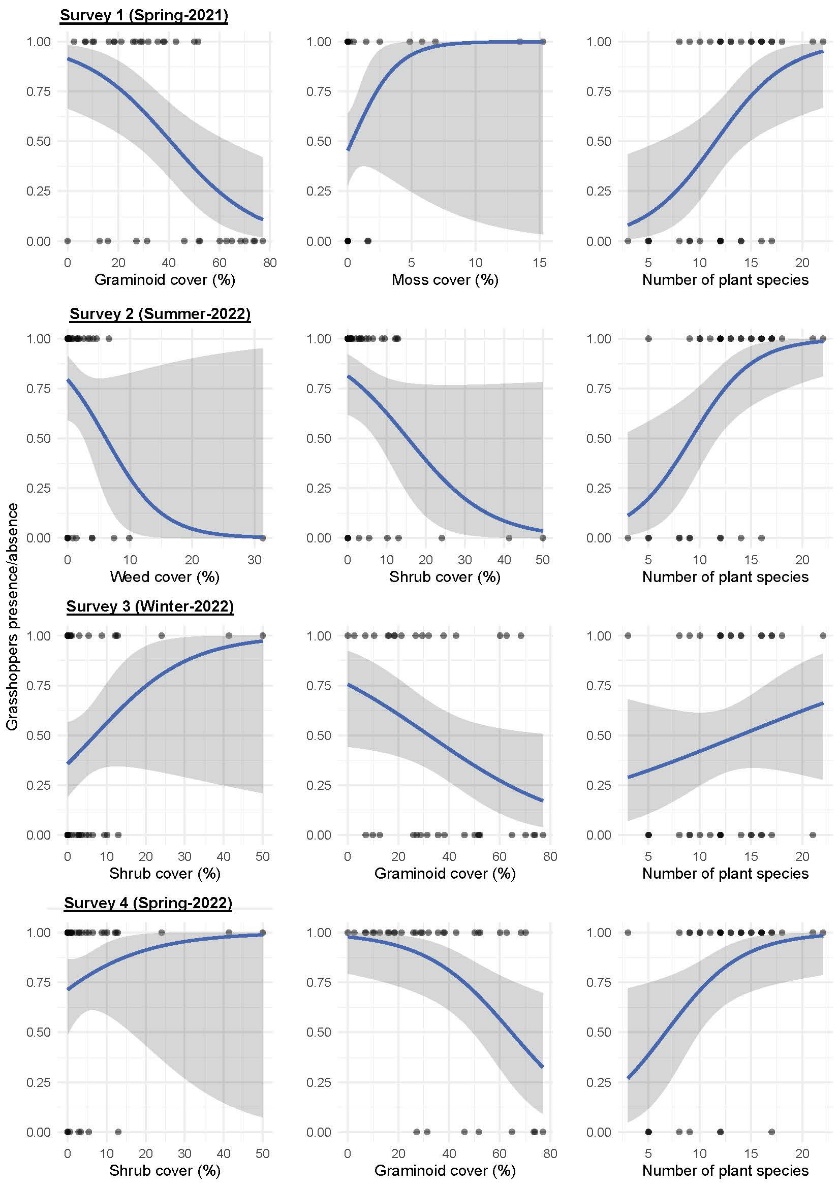

**Figure A3** a. Detection probability against soil temperature (standardized) and Occupancy probability against Graminoid cover, weed cover and plant species richness (Standardized). b. Number of individuals detected (Abundance) against habitat variables selected in the model with the highest Akaike weight per survey. The graphs were obtained fitting a negative-binomial model (estimated using ML). c. Presence (value=1) and absence (value=0) in relation to habitat variables selected in the model with the highest Akaike weight per survey. The relationships were obtained fitting a logistic model (estimated using ML)
